# MMS2plot: an R package for visualizing multiple MS/MS spectra for groups of modified and non-modified peptides

**DOI:** 10.1101/2020.03.13.989152

**Authors:** Liya Ming, Yang Zou, Yiming Zhao, Luna Zhang, Ningning He, Zhen Chen, Shawn S-C. Li, Lei Li

## Abstract

A large number of post-translational modifications (PTMs) in proteins are buried in the unassigned mass spectrometric (MS) spectra in shot-gun proteomics datasets. Because the modified peptide fragments are low in abundance relative to the corresponding non-modified versions, it is critical to develop tools that allow facile evaluation of assignment of PTMs based on the MS/MS spectra. Such tools would preferably have the ability to allow comparison of fragment ion spectra and retention time between the modified and unmodified peptide pairs or group. Herein, we describe MMS2plot, an R package for visualizing peptide-spectrum matches (PSMs) for multiple peptides. MMS2plot features a batch mode and generates the output images in vector graphics file format that facilitate evaluation and publication of the PSM assignment. We expect MMS2plot to play an important role in PTM discovery from large-scale proteomics datasets generated by LC (liquid chromatography)-MS/MS. The MMS2plot package is freely available at https://github.com/lileir/MMS2plot under the GPL-3 license.

## INTRODUCTION

The application of liquid chromatography-coupled tandem mass spectrometry (LC-MS/MS) in proteomics has generated an enormous amount of raw MS data, a large proportion of which have not been properly analyzed [1]. The main method used in MS-based protein identification is based on matching MS/MS spectra to peptide sequences and post-translational modifications (PTMs) contained in protein to PTM databases. However, with this approach, more than 50% of MS/MS spectra go unmatched, and many of the unmatched spectra represent peptides with unexpected modifications [2]. To detect these modifications, several unrestrictive methods for peptide identification have been proposed, including MSFragger and Open-pFind [2-6]. Furthermore, a number of approaches have been developed to determine the localization of the PTM based on results from the unrestrictive search [6, 7].

Nevertheless, the modified peptides to be identified are usually of low abundance and the corresponding MS/MS spectra are of poor quality. Probably due to this reason, the numbers of PTM PSMs identified by different programs [6, 7] are significantly different in the MS data of the human proteome. Therefore, it is necessary for the researchers to go through the tedious process of manually examining the MS/MS spectra one by one in order to validate the identification. This has significantly hindered the process of peptide and PTM identification. Consequently, the ability to easily visualize and compare raw MS/MS data is critical for data mining, spectrum quality monitoring and spectrum assignment for discovery proteomics.

The visualization of fragment ion spectrum annotation plays an important role in the validation of peptide identification. The annotated spectra can be visualized in three ways: 1) the single-spectrum view, 2) the mirrored-spectra view, which involves showing one spectrum above and the other below the x-axis for two spectra comparison, and 3) the aligned-spectra view, where more than two spectra are annotated by sharing the same x-axis. The visualization of mirror spectra or aligned spectra is preferable to inspect modified peptides in comparison to the corresponding non-modified peptides based on the following rationale. Modified peptides without enrichment are usually less abundant than their non-modified counterparts and the former is likely to be present in the sample only if the corresponding non-modified peptide has been detected.

Currently, several tools are available to annotate and visualize the PSMs. MS-Viewer, pFind, xiSPEC, PeptideShaker, PRIDE Inspector and IPSA provide a graphical user interface (GUI) for comprehensively annotating and visualizing multiple spectra [8-12]. However, they do not support either mirror or alignment plots. The integrative proteomics data viewer PDV supports a specific type of mirror plots in which two different spectra are matched to the same peptide or a single spectrum is assigned to two different peptides [13]. Msnbase and Spectrum_utils provide both single-spectrum view and mirrored-spectra view (Figure S1) [14, 15]. Nevertheless, both tools do not directly provide the sequence fragmentation plot where the peptide sequence is shown with separation of fragmentation keys, which is inconvenient for manual comparison between modified and non-modified peptides (See Figure S1 for details). Additionally, xiSPEC provides an annotated mirrored-spectra view (called Butterfly plot), yet this feature cannot be applied in an automated way useful for batch comparison of spectra [16]. In summary, the available tools support a single spectrum with the perfect annotation but few viewers are available to support both mirror and aligned spectra with comprehensive annotation.

Here we present the MMS2plot for displaying and comparing multiple PSMs assigned to non-modified peptides and the corresponding modified peptides in R. MMS2plot uses an automated analysis pipelines and offers both mirrored-spectra view and aligned-spectra view; and in either case, the displayed spectra share the same x-axis. The spectra view may be employed to compare mass shifts, intensities and matches of peaks between the modified and non-modified peptides. Additionally, it can be used to determine the shift in LC retention time due to changes in peptide hydrophobicity caused by the modified residue. Compared with the available tools, MMS2plot provides several unique features: 1) it can display multiple PSMs per window; 2) the displayed PSMs can be modified/annotated by the user, which is important for the PTM localization; 3) the data can be output in batch mode in vector graphics file formats. MMS2plot is freely available as open source at https://github.com/lileir/MMS2plot.

## Methods & results

MMS2plot was based on several R packages such as Msnbase and mzID for speeding up many processing steps. MMS2plot requires five different input files, as follows (Figure 1):

**Figure 1.**
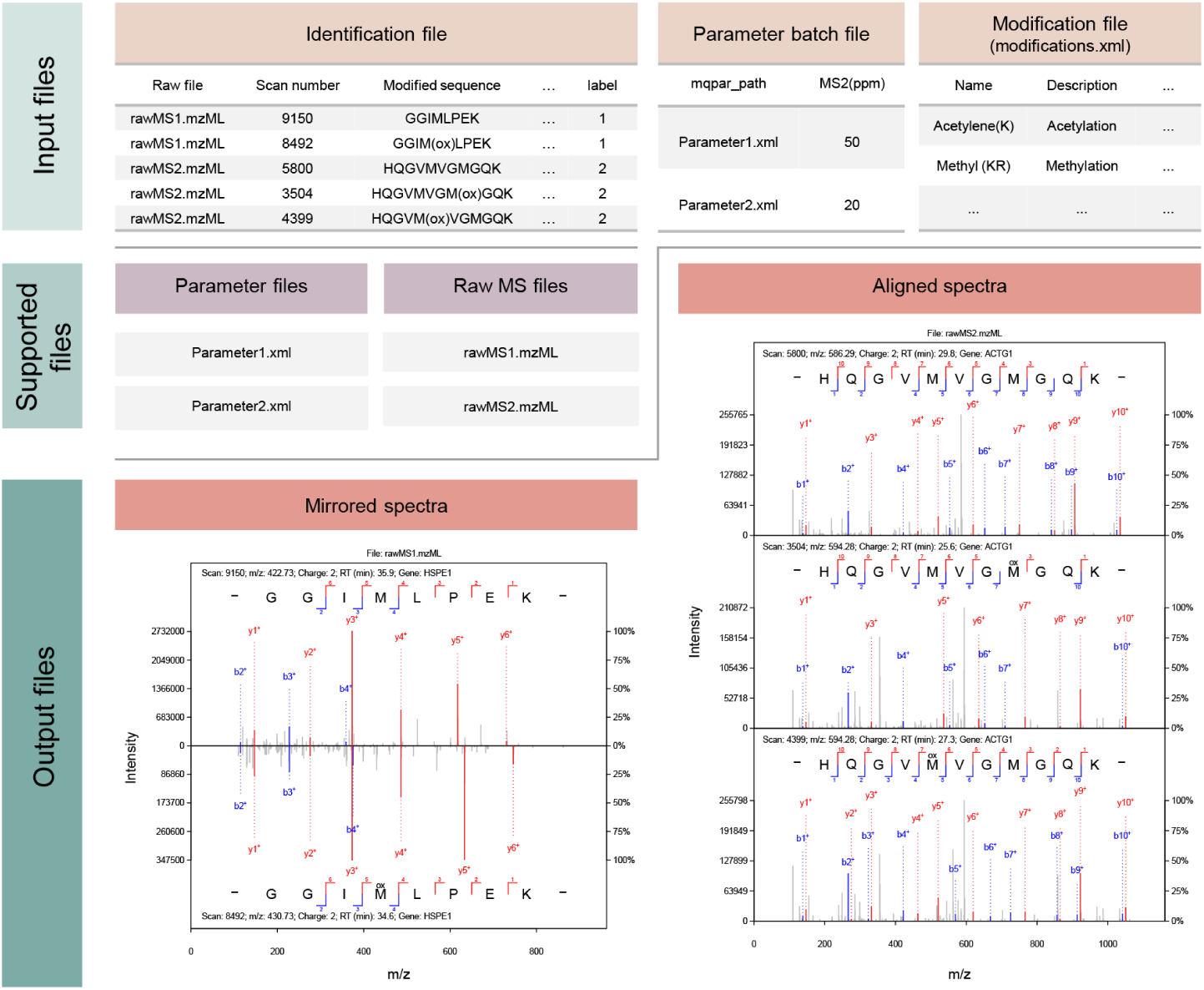
An overview of MMS2plot. The input files for MMS2plot and the related supported files are shown at the top. Examples of output images are shown at the bottom.

1. A modification configuration XML file where the types of protein modifications are defined. It has the same XML format as the “modifications.xml” file in MaxQuant [8]. Thus, user-defined modifications can be configured in MaxQuant and the resulting modifications.xml file can then be used directly in MMS2plot.
2. A parameter XML file that includes the raw MS filenames and the modifications that are defined for MS spectral analysis. This parameter file has the same XML format as the mqpar.xml file in MaxQuant. Similar to the modification file described above, the mqpar.xml file can be directly imported from MaxQuant to MMS2plot.
3. A parameter batch text file that contains the parameter XML files to be processed, the corresponding settings in fragment mass tolerance and the ion types to be shown (i.e. b/c/y/c ions). The parameter files should be provided with the full file path and the mass tolerance should be set in the Parts per Million (PPM).
4. An identification text file that contains the PSM information of all peptides to be displayed and their group number. The PSM information can derived from the results of MaxQuant, pFind, X!Tandem and other search engines. Other identification formats can be converted to this text format using the related R packages. For instance, the mzIdentML format could be converted using the mzID package (see the user guide for the details)[17]. In this file, each row lists a single PSM plus the group number assigned to the PSM by the user. The PSMs with the same group label will share the same x-axis and be output together. In other words, distinct spectra assigned to different peptides can be shown together if they are assigned to the same group number. The advantage of the text format is that the PTM location is easily modified by the user for manual inspection.
5. The raw MS files in standard mzML format. The conversion of MS data files to the mzML format can be done using the MSConvert application tool in ProteoWizard [18].

Please note that the two XML files described above are related to MaxQuant configuration files and they are easy to generate. MMS2plot supports the results from any search engine rather than MaxQaunt. Nevertheless, these results are required to be converted into the text format using existing functions from R or other computer languages. Based on these inputs, MMS2plot automatically loads both parameter files and the raw MS files, extracts the MS2 spectra, calculates the PSMs using user-defined fragment mass tolerance, and generate images of the spectra. MMS2plot supports both label-free data and labelled data, including those from TMT, SILAC, and dimethyl-labelling (See examples in the MMS2plot package). MMS2plot not only support neutral loss specific to a PTM type such as phosphorylation (Figure S2A) but also different types of ions (Figure S2B). When a fragment ion can be matched to multiple ions of a peptide, all the matched ions will be labelled and shown (Figure S3). If the parent ion charge is three or more, fragment ion charge as one or two will be considered only. The output images display both the absolute and relative intensities of the spectra, the sequences assigned to the spectra with the type of PTM shown on top of the modified residue, the raw MS files from which the spectra are derived, the spectra and additional information (e.g., number of scans, m/z, charge, retention time, and gene names of the corresponding proteins where the matched peptides are derived). MMS2plot allows users to adjust the spectral image such as the size of the image, the color of (un)matched peaks, the margin of the spectral figure for both the x and y axes (Figure S4). MMS2plot comes with a default set of parameters to facilitate visualization and publication of the PSM data. By default, the width of the image is preset at 3.35 inch for a single column and 7 inch for double columns.

We used a few examples to demonstrate the merits of MMS2plot. Figure 2A shows that compared with the spectrum (Scan number: 10774) annotated with the unmodified peptide “GGGGNFGPGPGSNFR”, the spectrum (Scan number: 10874) in the same MS raw file is a mixture spectrum probably corresponding to the same peptide sequence with different types, where the peptide is oxidized at P10 or F6 [19]. This spectrum may also correspond to the substitution of F6 with Y6 [20]. Clearly, the aligned-spectra view shows more information than the single-spectrum view. Figure 2B shows that K56 at CHID1 (Chitinase domain-containing protein 1) can be hydroxylated and this modified site is the cleavage site. The related miscleaved modified and non-modified peptides were also found in the same raw MS file (Figure S5). These indicate that the CHID1 K56 is a novel hydroxylated site and hydroxylated lysine can be digested by trypsin, consistent with the previous observation [21]. In both cases, the fragment ions in the modified peptides are of lower abundance than those in the corresponding non-modified peptides and both have the similar retention time. Figure S3 shows that the annotation of the upper spectrum in the mirror spectra as the modified peptide is probably wrong as the matched ion peaks are significantly different from those annotated to the unmodified counterpart in the bottom spectrum. More information including how to install this package, how to run the examples in the help(mms2plot) page and how to generate MMS2plot required files was described in the user guide as Supplementary file 1. A packed example of MM2plot required files is accessible at https://github.com/lileir/MMS2plot.

**Figure 2.**
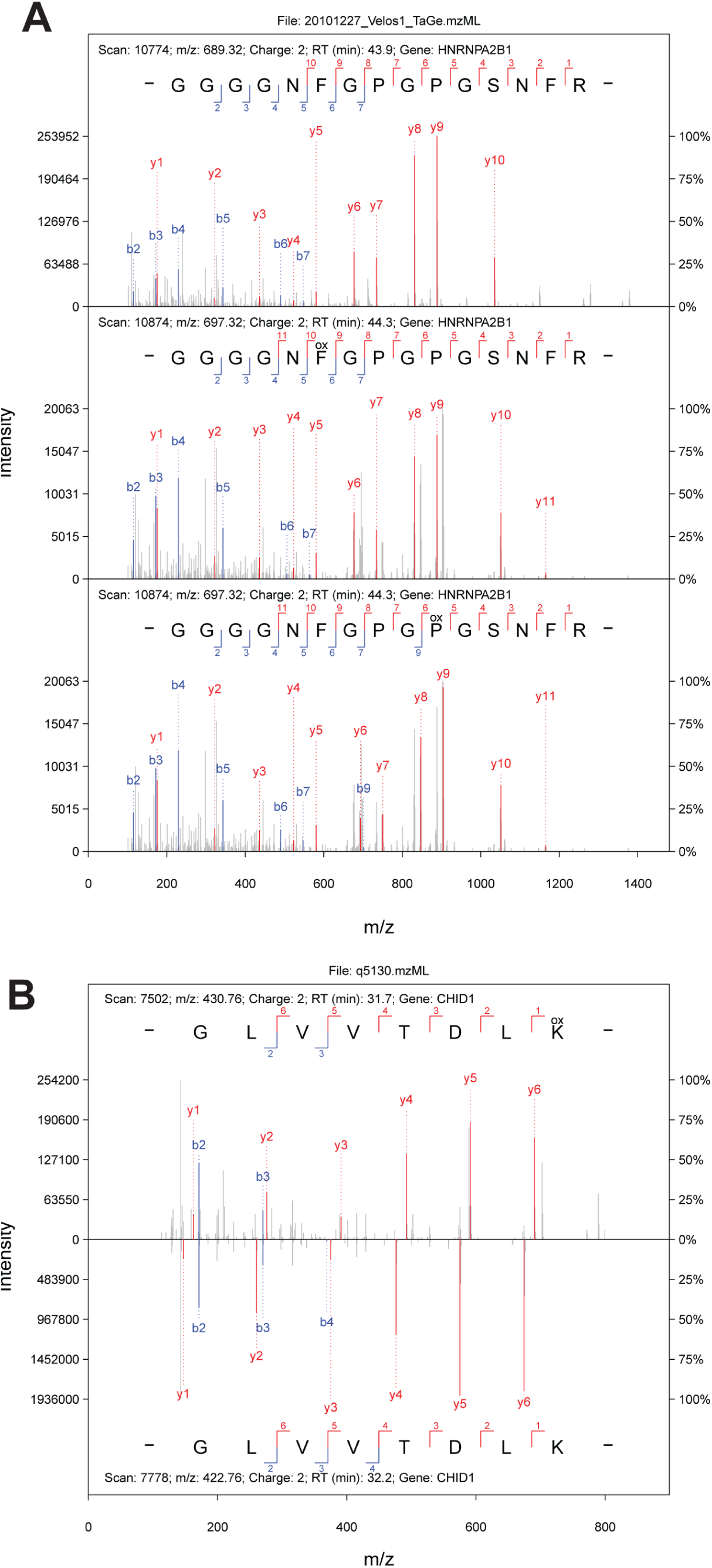
Two examples of MMS2plot. **A**. An aligned spectra of novel oxidized peptides with the unmodified counterpart. The spectrum containing the oxidation is probably a mixed spectrum, including both proline-oxidized peptides and phenylalanine-oxidized peptide. **B**. A mirror spectra of a novel lysine oxidized peptide with the unmodified counterpart.

## Discussion

A number of approaches have been developed to date to allow analysis of unassigned spectra in MS/MS datasets generated from shotgun proteomics. Some of the spectra are found to represent modified peptides [6] and others represent peptides with amino acid substitutions [22]. Although thousands of such peptides have been discovered, a suitable tool is lacking that allows visual display of the corresponding PSMs for peak assignment and validation of identification. To address this shortcoming in proteomics analysis, we have developed the MMS2plot software that can align and display multiple related spectra in a single image, thereby allowing the user to compare the spectra and evaluate assignment. The output images are in the vector graphics file format with preset configurations suitable for inclusion as figures in publication. One possible disadvantage of MMS2plot is that a graphical user interface is not provided for visualization and to help users adjust the settings and produce ideal images for publication. We reason that because MMS2plot is designed to produce a large number of images simultaneously, it is impractical to manually adjust settings to fit all the images. Alternatively, users may adjust the settings for specific annotated spectra according to the outputted images or preset the settings based on the visual effects using other visualization tools. Indeed, IPSA provided GUI for the annotation of a single spectrum but not for bulk processing of spectra annotation [8].

## Acknowledgments

This work was supported, in part, by grants (to SSCL) from the Canadian Institute of Health Research (CIHR), funds from the Young Scientists Fund of the National Natural Science Foundation of China (Grant No. 31701142 to ZC; Grant No. 81602621 to NH), the Shandong Provincial Natural Science Foundation (Grant No. ZR2016CM14 to LL), the National Natural Science Foundation of China (Grant No. 31770821 to LL); LL is also supported by the “Distinguished Expert of Overseas Tai Shan Scholar” program. SSCL held a Canada Research Chair (Tier I) in Molecular and Epigenetic Basis of Cancer.

## Authors’ contributions

LL and SSCL conceived and designed the project. LM, YZ YMZ and ZC developed the algorithms; LM, NH and LNZ provided examples; LL, SSCL and LM wrote the manuscript. All authors read and approved the final manuscript.

