## Supplementary Figures for "MMS2plot: an R package for visualizing multiple MS/MS spectra for groups of modified and non-modified peptides"

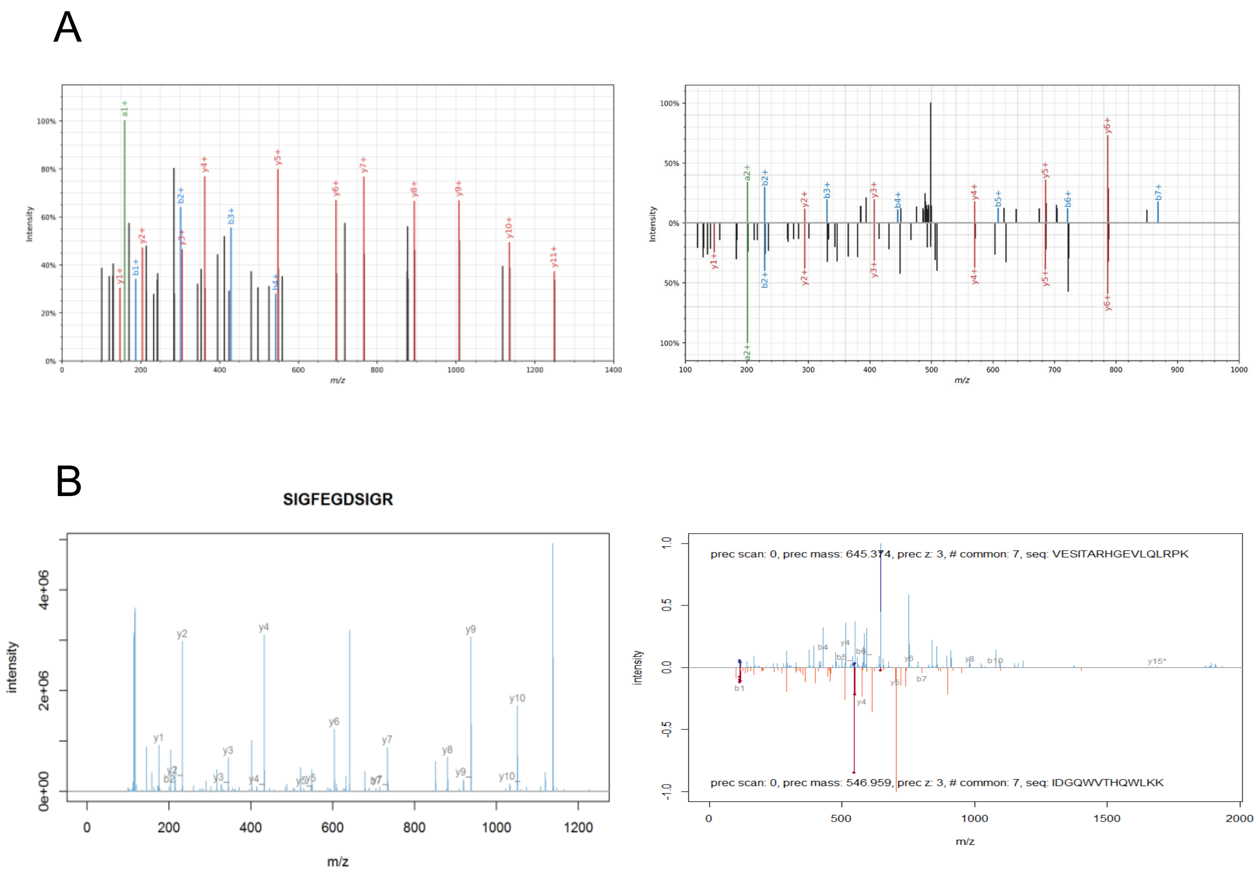


**Figure S1.** Examples of annotated spectra extracted from spectrum_utils [1] (**A**) and MSnbase [2] (**B**), where the left shows a single spectrum and the right shows a mirrored spectra.

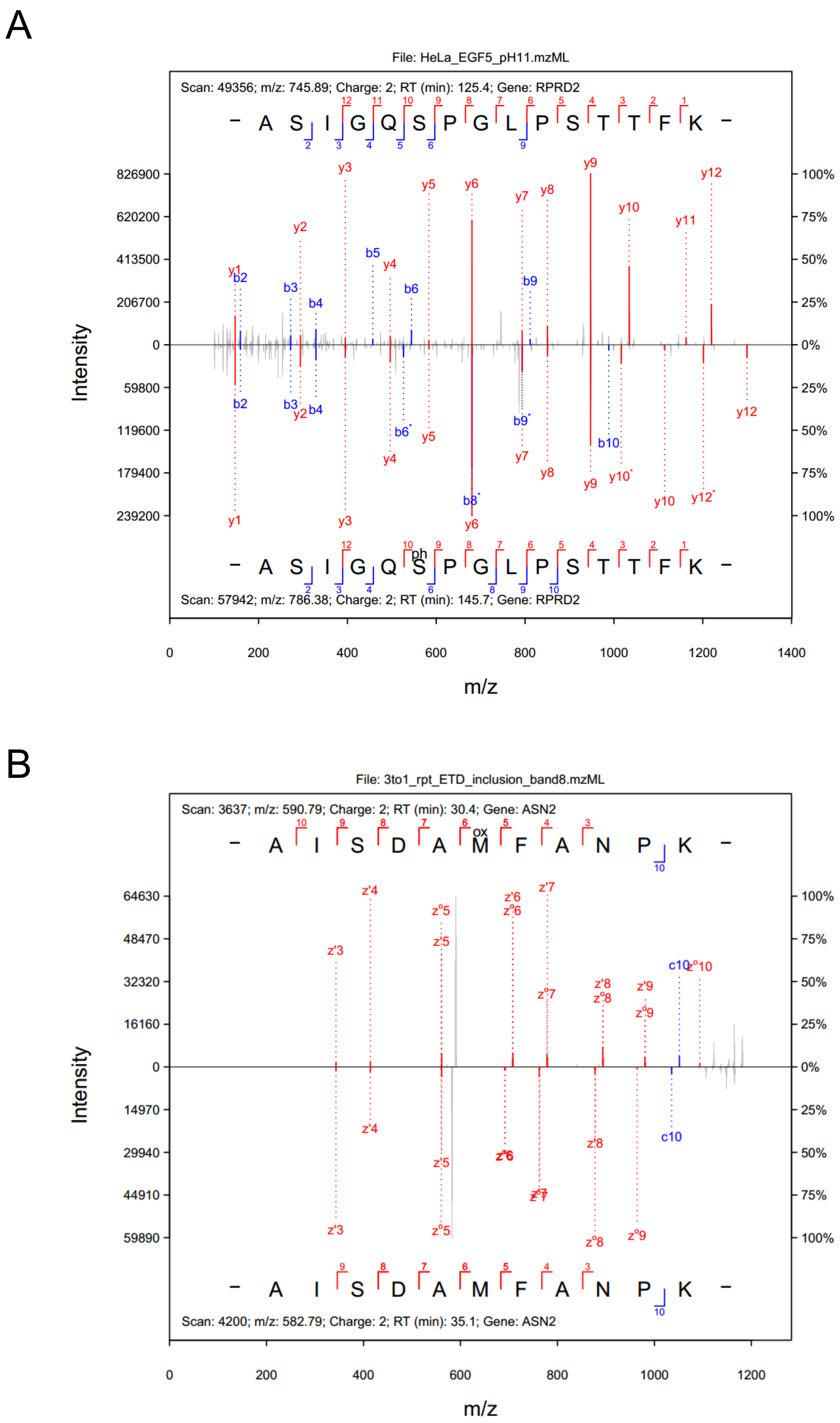


**Figure S2.** MMS2plot examples of spectra annotated with neutral loss (marked as *) (**A**), annotated with c/z ions (**B**).

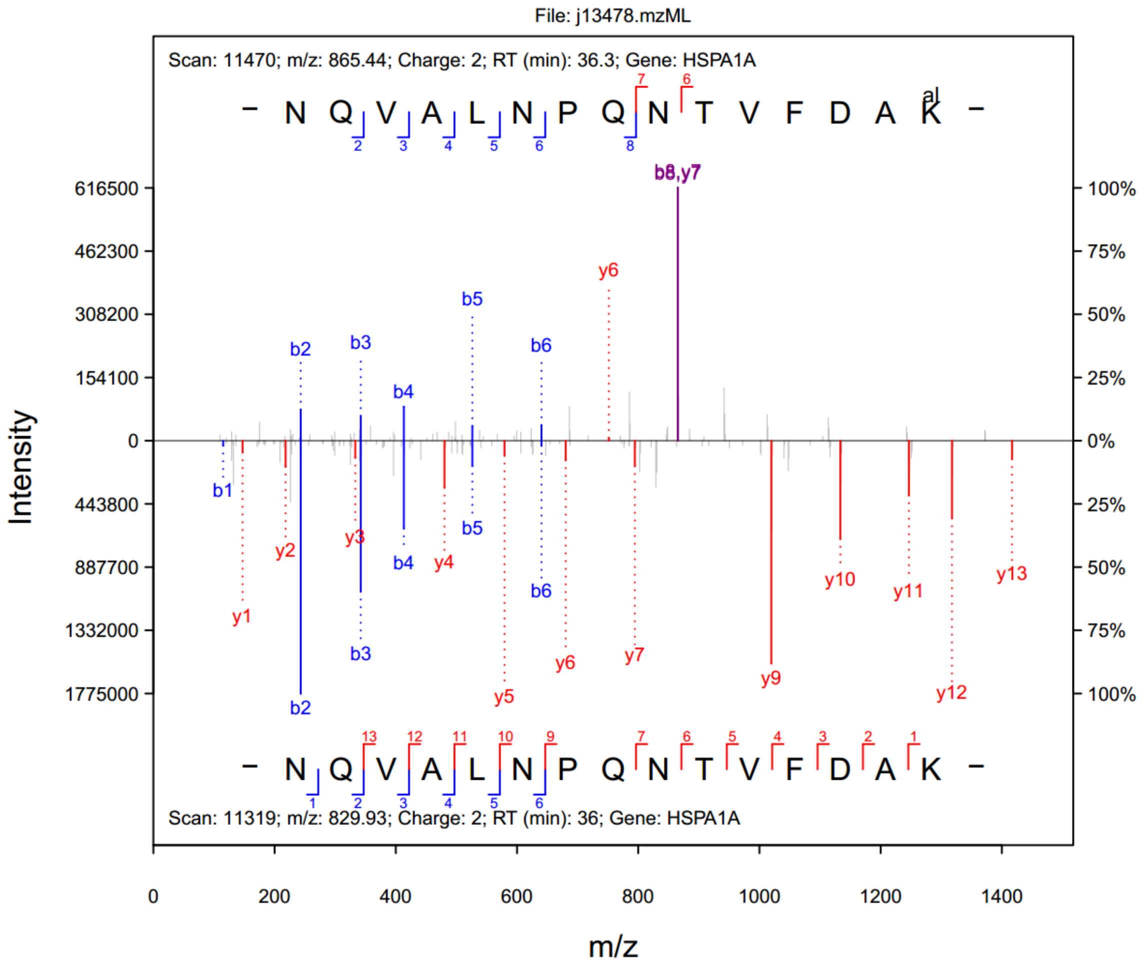


**Figure S3.** A mirror spectra of a novel lysine alaninylated peptide with the unmodified counterpart. Lysine aminoacylation has recently been discovered that has biological functions [1]. Both b8 and y7 ions are assigned to the same fragment ion which can be matched to both ions. The significant difference between the matched ion patterns for the upper spectrum and the bottom spectrum indicates that the annotation of the upper spectrum as the modified peptide is probably incorrect.

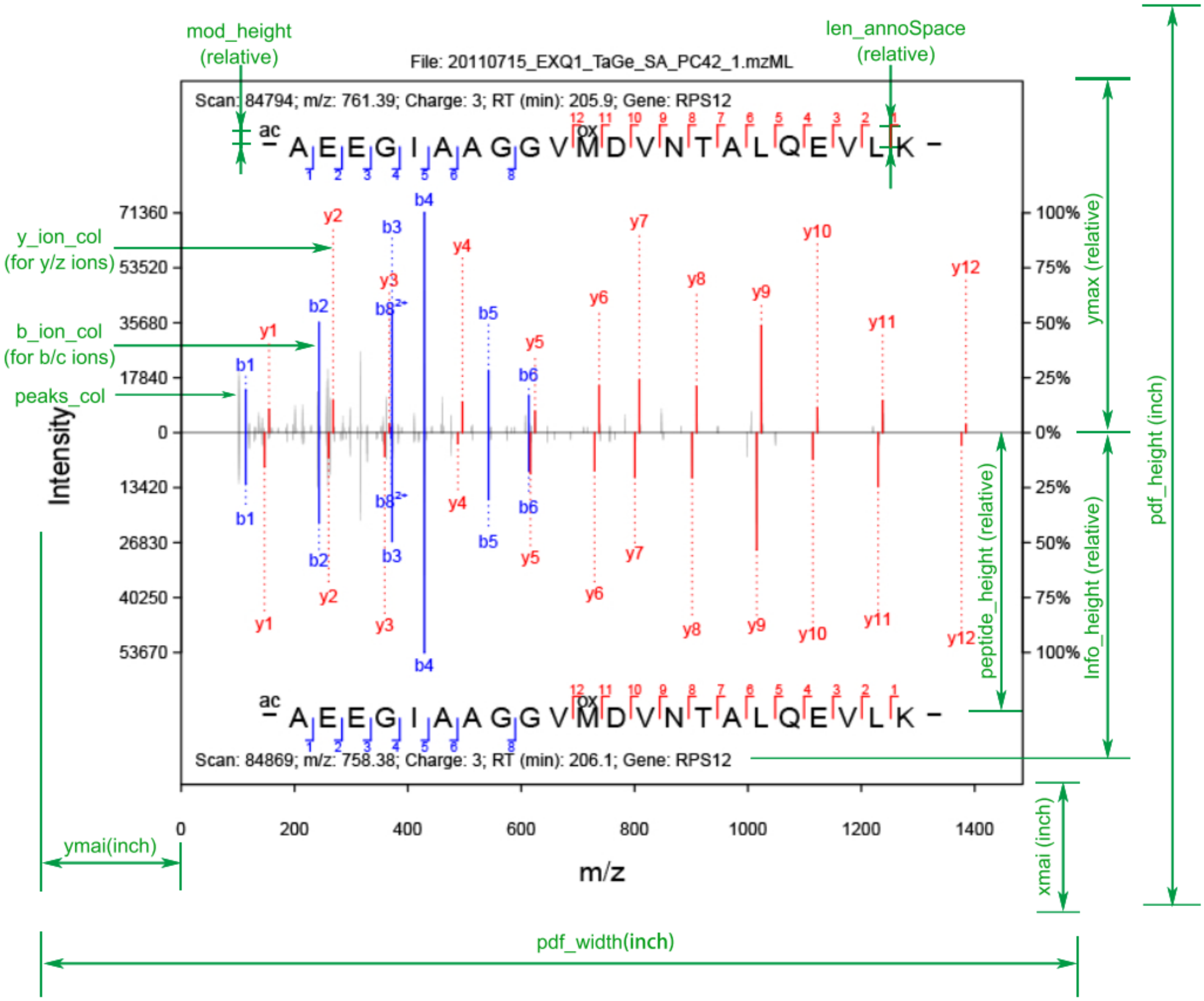


**Figure S4.** The parameters that users can set to adjust the output images in MMS2plot.

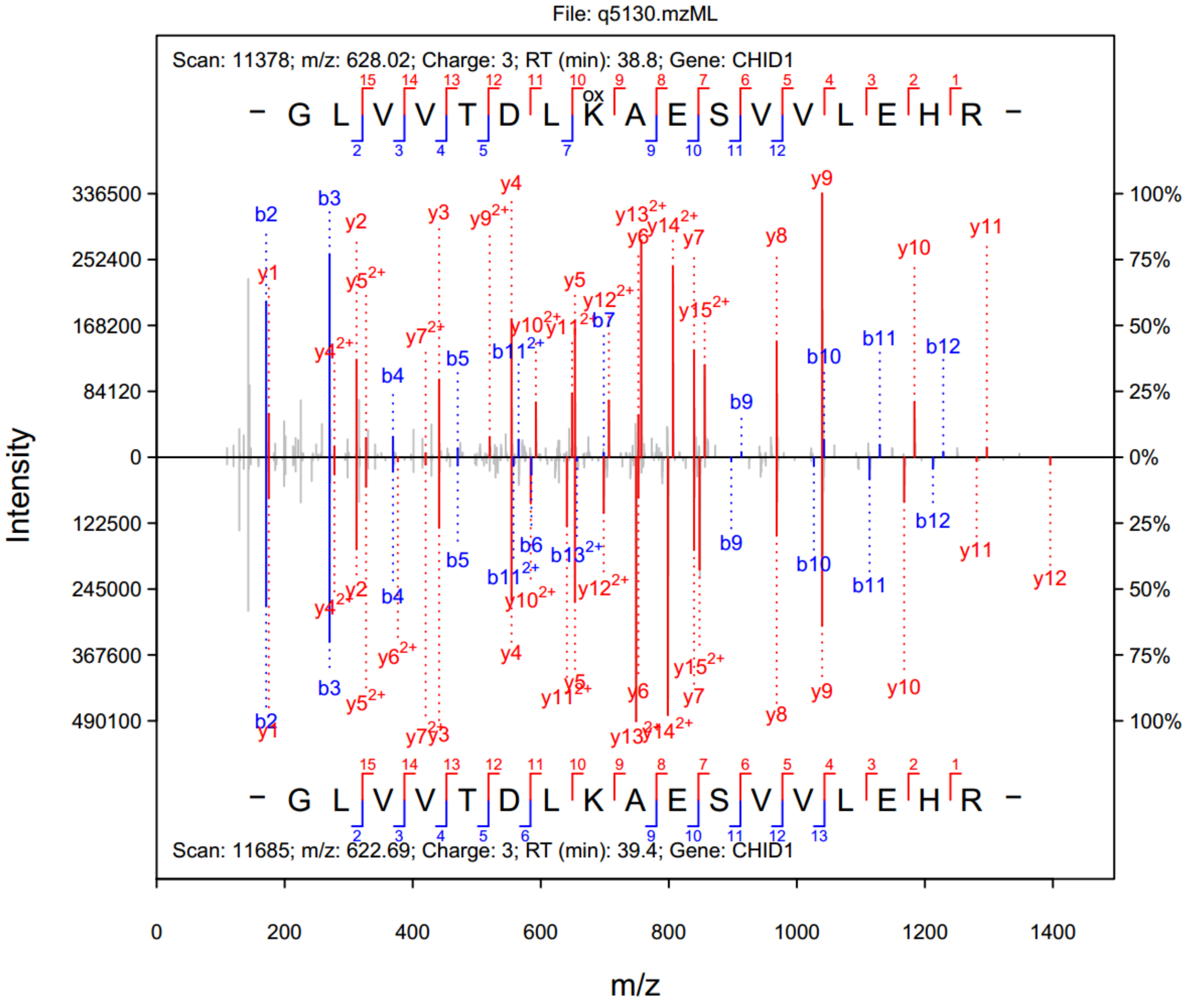


**Figure S5.** Miscleaved hydroxylated lysine-containing peptide and non-modified counterpart were found in the raw MS file q5130.mzML.
