## Supplementary file for "MMS2plot: an R package for visualizing multiple MS/MS spectra for groups of modified and non-modified peptides"

### MMS2plot user guide

### (version 1.01.09)

Written on January 7, 2020

#### Brief introduction

MMS2plot, an R package for visualizing and comparing peptide-spectrum matches (PSMs) assigned to non-modified peptides and the corresponding modified peptides, compatible with proteomics data generated from LC-MS/MS analysis. MMS2plot uses an automated analysis pipelines and offers both mirrored-spectra view and aligned-spectra view; and in either case, the displayed spectra share the same x-axis. The spectra view may be employed to compare mass shifts, intensities and matches of peaks between the modified and non-modified peptides. Additionally, MMS2plot features a batch mode and generates the output images in vector graphics file format that facilitate evaluation and publication of the PSM assignment.

### 1. Installation of MMS2plot

The MMS2plot R package supports on multi-OS systems (such as Windows and Linux) and is freely available through https://github.com/lileir/MMS2plot. It can be directly installed from GitHub using the library "devtools" under the R environment.

Use the following command to install and load MMS2plot:

```
>if(!requireNamespace("devtools"))
install.packages("devtools")
>library(devtools)
>devtools::install_github("lileir/mms2plot")
>library(mms2plot)
```

Note:

1. The version of the "devtools" package should be >2.2. Otherwise, the "mms2plot" package cannot be installed successfully.
2. The MMS2plot package depends on R (>= 3.5) and requires the following packages (e.g. xml2, data.table, DescTools, MSnbase, scales, Biobase, gsubfn, graphics and grDevices).
3. The failure of the "mms2plot" installation is mainly due to the following reasons:
   - The required packages previously installed are out of date, especially for the "devtools" package. Please update the packages.
   - The error message is usually shown as : "Error in utils::download.file(url, path, method = download_method(), quiet = quiet, : cannot open URL 'https://api.github.com/repos/lileir/mms2plot/tarball/master'". This failure is due to the slow internet speed. Please re-run the command above.
   - Failure of installation of the required packages, especially for iOS or linux. We suggest installing the binary version of the required packages. If the error "'tar: Failed to set default locale' error?" appears, please refer to the link https://stackoverflow.com/questions/3907719/how-to-fix-tar-failed-to-set-def

ault-locale-error/45469967#45469967. If you prefer their source versions, please make sure that the compilation environment works fine.

4. After installation and loading of this package, the command "help(mms2plot)" can be executed to view the help information.

### 2. Examples in the help(mms2plot) page

After the installation of the MMS2plot package, the typical thing a user would like to do is to follow the example in the help page. Figure 1 shows the commands and the output of the annotated spectra from the label-free data.

A

```
> library(mms2plot)
> general_path = system.file( package = "mms2plot",dir = "extdata" )
> setwd( general_path )
> # Generate a mirrored-spectra pdf file for label-free data
> lf_path = system.file( package = "mms2plot",dir = "extdata/label_free" )
> # id_table_path expands the Maxqaunt output msms.txt by adding "label" column
> id_table_path = dir( lf_path, "msms_labelfree.txt", full.names = TRUE )
> mod_xml_path = dir( general_path, "modifications.xml", full.names = TRUE )
> # par_filepath contains par.xml with full file path and PPM cutoff
> par_filepath = dir( general_path, "par_batch.txt", full.names = TRUE )
> output_path = general_path
> mms2plot( id_table_path, mod_xml_path, par_filepath, output_path) # Only run for testing

[1] "Reading the raw MS file:  label_free/j6577.mzML ... ..."
[1] "Reading ... ... completed!"
[1] "The pdf file 'C:/Program Files/R/R-
3.6.2/library/mms2plot/extdata/j6577.mzML_1_AALEKPMDLEEEKK_mirror.pdf' was generated."
```

B

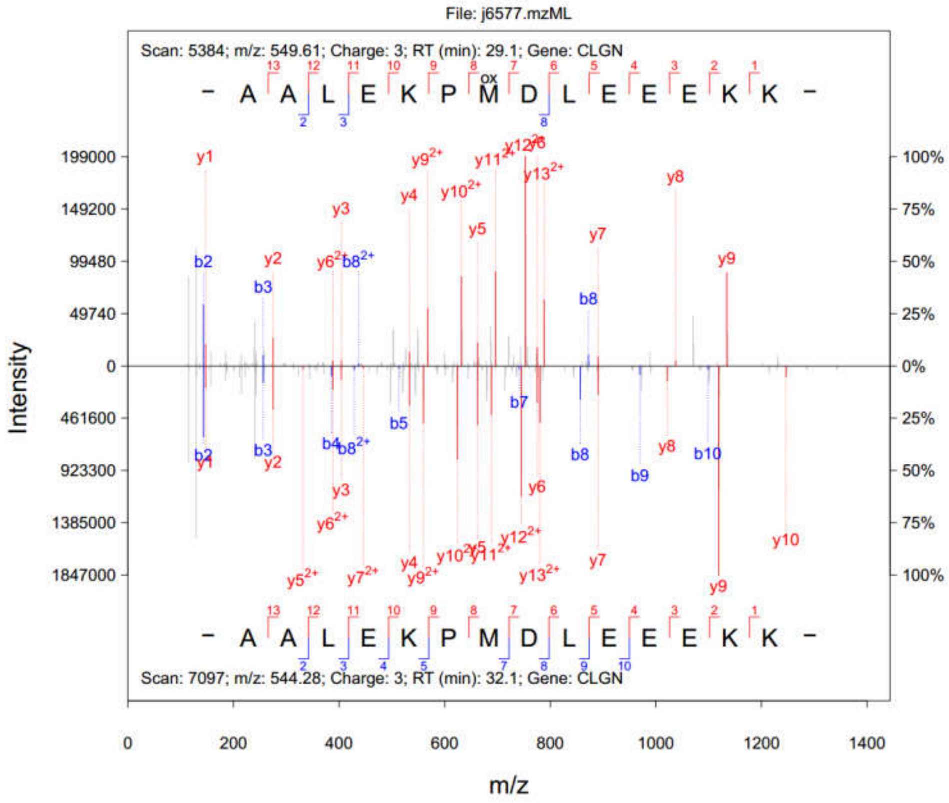

**Figure 1.** The commands (**A**) and the output (**B**) of for the label-free example in the mms2plot help page.

### 3. Generation of MMS2plot required files

The MMS2plot required files include both input files and supported files. **Supplementary Data 1** contains the example files. It can be downloaded and unzipped to a given folder (e.g. "D:\test"). This folder includes two subfolders:

1) "input", which covers three files: modifications.xml, identification.txt and parameter_batch.txt;
2) "support", including two parameter files (parameter1.xml & parameter2.xml) and two raw MS files (rawMS1.mzML & rawMS2.mzML).

These required files are described as below:

#### 3.1 The modifications.xml file

This file is directly derived from the Maxquant software. If the modification of interest is not included in the xml file, this modification can be added to the xml file through Maxquant. In the some versions of Maxquant, the modification information is stored in two xml files: modifications.xml and modifications.local.xml. Please combine them as a single xml file, by pasting the content of the second file to the end of the first file, followed by removing the last line of "</modifications>" from the first file and the first two lines from the second file.

#### 3.2 The identification.txt file

This text file contains eight necessary columns (as listed below) whereas any other columns will be ignored (**Figure 2**).

1) "Raw file": the full file path of the MS raw file with the mzML format, for example, "D:\test\support\rawMS1.mzML" for Windows OS and "/test/support/rawMS1.mzML" for Linux OS.
2) "Scan number": the RAW-file derived scan number of the MS/MS.
3) "Sequence": the identified Amino acid sequence of the peptide.
4) "Modifications": PTM types contained within the sequence, for instance, "Oxidation(M)". When no modifications exist, "Unmodified" should be filled.

5) “Modified sequence”, sequence representation of the peptide including location(s) of modified amino acids.

6) “Gene Names”: names of genes that the identified peptide is associated with.

7) “Charge”: the charge state of the precursor ion.

8) “label”: a number assigned to the peptide-spectra matches (PSMs) that share the x-axis (*i.e.* m/z) of the output image.

| Raw file | Scan number | Sequence | Modifications | Modified sequence | Gene Names | Charge | label |
|---|---|---|---|---|---|---|---|
| D:\test\support\rawMS1.mzML | 9150 | GGIMLPEK | Unmodified | GGIMLPEK | HSPE1 | 2 | 1 |
| D:\test\support\rawMS1.mzML | 8492 | GGIMLPEK | Oxidation(M) | GGIM(ox)LPEK | HSPE1 | 2 | 1 |
| D:\test\support\rawMS2.mzML | 5800 | HQGVMVGMGQK | Unmodified | HQGVMVGMGQK | ACTG1 | 2 | 2 |
| D:\test\support\rawMS2.mzML | 3504 | HQGVMVGMGQK | Oxidation(M) | HQGVMVGM(ox)GQK | ACTG1 | 2 | 2 |
| D:\test\support\rawMS2.mzML | 4399 | HQGVMVGMGQK | Oxidation(M) | HQGVM(ox)VGMGQK | ACTG1 | 2 | 2 |

**Figure 2**. An example of the identification.txt file that contains the information of five PSMs. The first two are used to generate the mirrored spectra and the other three are employed to produce the aligned spectra.

##### 3.3 Conversion of the identification file from the mzidentML file format

The identification text file is easily generated from the Maxquant search result. Other identification file formats are easily converted to this text file format. For instance, the mzIdentML format can be easily converted to the text file format using two functions (i.e. mzID and flatten) of the mzID package, as an mzIdentML parser for R (**Figure 3**). After the conversion, users need to further modify the output text file so that it could be used as the MMS2plot input file (**Figure 3**). For example, users need to add the “label” column for grouping the annotated spectra. Please note that the annotated peptides as the MMS2plot input is based on, but does not rely on, the identifications from the search engines. User can easily modify the PTM location or even the whole peptide for manual inspection.

We described the detailed process by using the following example. We used the search engine MSGFplus to identify the oxidized peptides from the raw file q5145.raw (MSGFplus usage can be found using the command “java -Xmx3500M -jar MSGFPlus.jar”). We converted the raw file to the standard mzML file format using MSConvert as the mzML file is required as an input of MSGFplus.

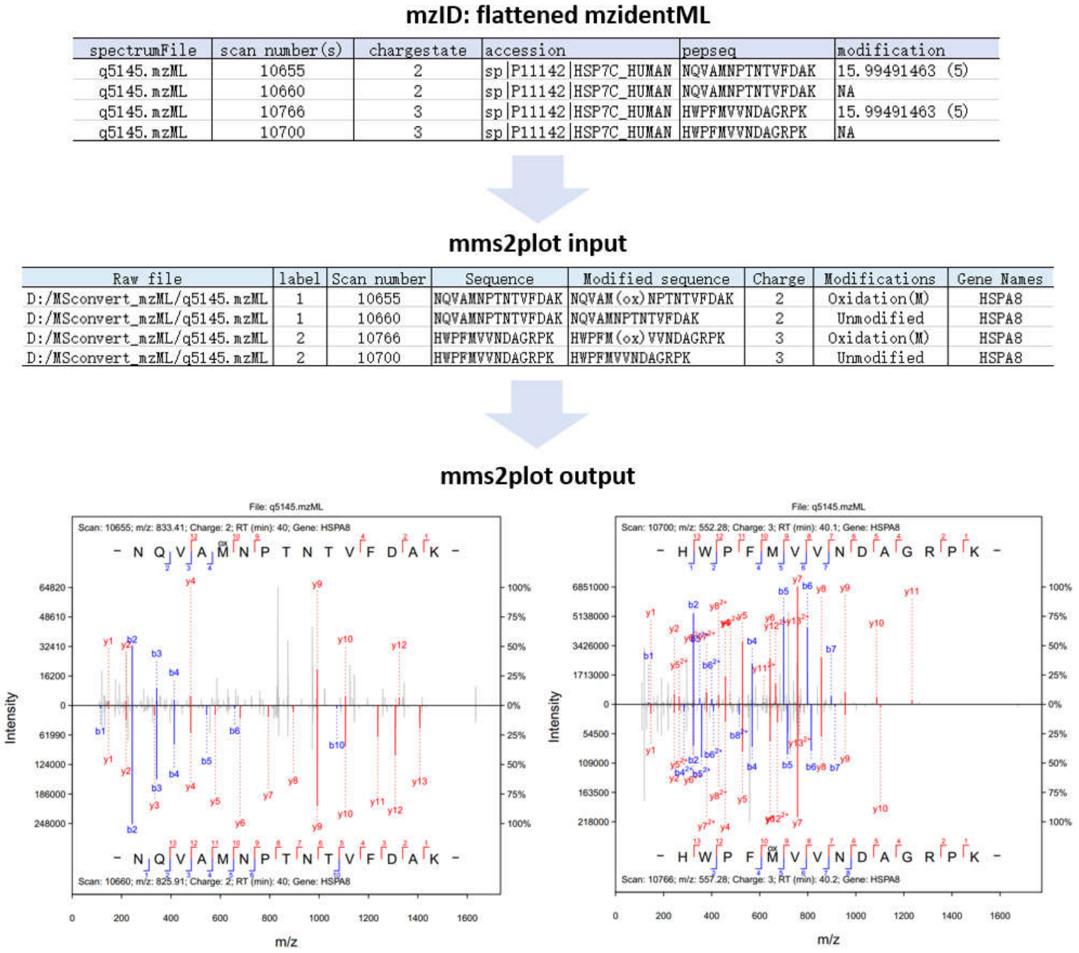

**Figure 3**. The procedure from conversion of the mzIDentML file format to the input file of MMS2plot to the MMS2plot. The related commands are listed below.

We then got the identification file with the default mzIdentML file format (i.e. q5145.mzid) via the following command:

```
> java -Xmx3500M -jar MSGFPlus.jar -s q5145.mzML -d test.fasta
-mod Mods.txt -o q5145.mzid
```

The running process was shown below.

```
Search Parameters:
	PrecursorMassTolerance: 20.0ppm
	IsotopeError: 0,1
	TargetDecoyAnalysis: false
	FragmentationMethod: As written in the spectrum or CID if no info
	Instrument: LowRes
	Enzyme: Tryp
	Protocol: Standard
	NumTolerableTermini: 2
	MinPeptideLength: 6
	MaxPeptideLength: 40
	NumMatchesPerSpec: 1
Spectrum 0-907 (total: 908)
pool-1-thread-1: Preprocessing spectra...
Loading built-in param file: HCD_QExactive_Tryp.param
pool-1-thread-1: Preprocessing spectra finished (elapsed time: 3.00 sec)
pool-1-thread-1: Database search...
pool-1-thread-1: Database search progress... 0.0% complete
pool-1-thread-1: Database search finished (elapsed time: 0.00 sec)
pool-1-thread-1: Computing spectral E-values...
pool-1-thread-1: Computing spectral E-values finished (elapsed time: 4.00 sec)
Writing results...
Writing results finished (elapsed time: 0.51 sec)
MS-GF+ complete (total elapsed time: 10.28 sec)
```

The generated mzidentML file could be flattened as the text file using the following commands:

```
>if (!requireNamespace("BiocManager", quietly = TRUE))
install.packages("BiocManager")

>BiocManager::install("mzID")

>library(mzID)

>path_mzid <- "D:/MSGFplus/q5145.mzid"  #path of the
mzidentML file

>mzidResult <- mzID::mzID(path_mzid)

>flatResults <- mzID::flatten(mzidResult)
```

The result from the function ‘mzID::flatten’ contains multiple columns, out of which six are required for MMS2plot (**Figure 3**).

#### 3.4 The parameter batch text file

This text file (named as parameter_batch.txt in **supplementary data 1**) contains three columns (**Figure 4**):

1) the full file path of the parameter XML files (see details in **3.5**);

2) the fragment mass tolerance set in the Parts per Million (ppm) for each XML file;

3) the ion types to be shown (*i.e.* y, z and yz, where y stands for b/y ions, z stands for c/z ions and yz stands for b/c/y/z ions, respectively).

| par_path | ppm | ion_type |
|---|---|---|
| D:\support_folder\parameter1.xml | 20 | y |
| D:\support_folder\parameter2.xml | 20 | y |

**Figure 4**. An example of the parameter batch file that contains two parameter files.

#### 3.5 The parameter XML file

This file includes the raw MS filenames and the modifications that are defined for MS spectral analysis. This parameter file is the same as the mqpar.xml file in Maxquant and thus it can be directly generated using Maxquant.

#### 3.6 Conversion of the mzML files from the raw MS files

MMS2plot only support the standard mzML file format as the raw MS data input. Take the two mzML files shown in **Figure 2** as example. The conversion of MS data files to the mzML format can be done using the MSConvert application tool in ProteoWizard. In addition to the **msConvertGUI** program, the command line tool **msconvert** is more efficient for batch conversion.

```
"C:\ProteoWizard\msconvert.exe" D:\Raw\rawMS1.raw --filter
"scanNumber [9150,9150] [8492,8492]" -o D:\test\support
```

As a result, two MS2s (scan number 9150 and 8492) from rawMS1.raw was extracted as the mzML file format and saved to the designated path.

### 4. Argument settings of MMS2plot

The function 'mms2plot' requires four necessary input arguments while the other arguments have default values.

The four arguments necessary represent the following respectively:

1) "id_table_path", the path of the identification file ;
2) "mod_xml_path", the path of the modification xml file;
3) "par_filepath", the path of the parameter batch file;
4) "output_path", the path of the folder used to save the output spectra.

For example:

```
>id_table_path <- "D:/test/input/identification.txt"
>mod_xml_path <- "D: /test/input/modifications.xml"
>par_filepath <- "D:/test/input/parameter_batch.txt"
>output_path <- "D:/test"
>mms2plot(id_table_path, mod_xml_path, par_filepath,
output_path)
```

Note: In windows OS, a single forward slash (or double backslashes) is used to separate the folders.

### 5. MMS2plot output

The pdf files were generated and saved to the specified path (e.g. “D:\test”). The pdf file name is designated to contain four types of information, separated by the underline characters (**Figure 5&6**). The information includes 1) the raw MS file name, 2) the label number, 3) the identified peptide sequence and 4) the type of visualization (mirror or align).

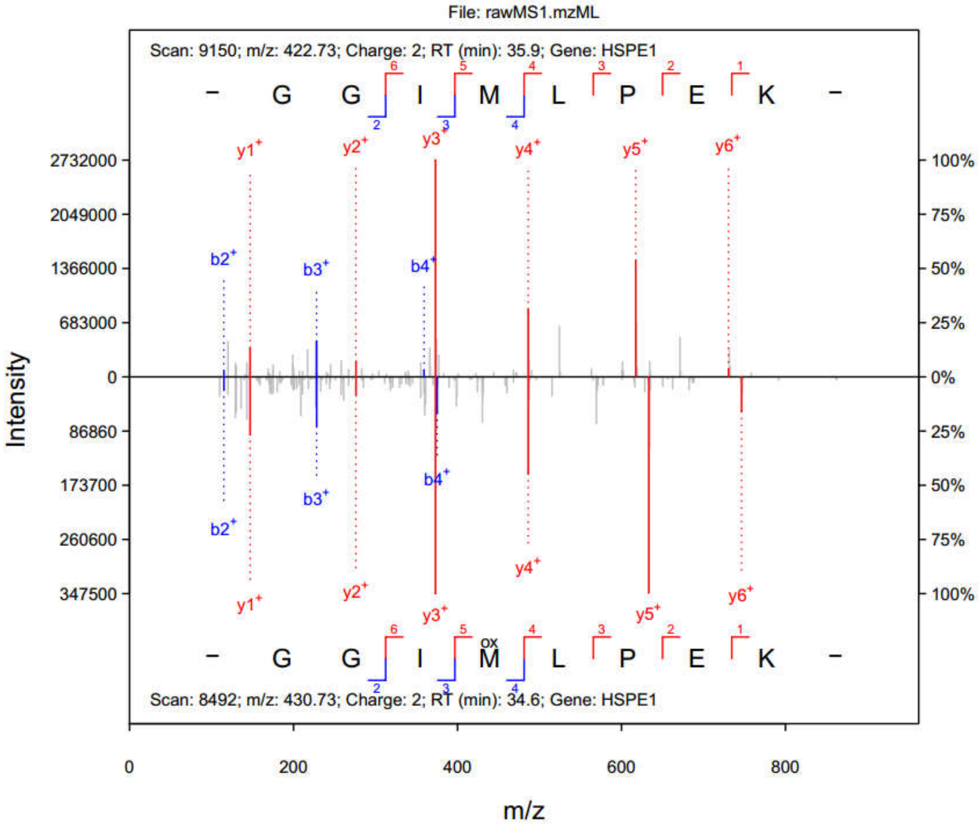

**Figure 5**. The image of the file “rawMS1.mzML_1_GGIMLPEK_mirror.pdf”.

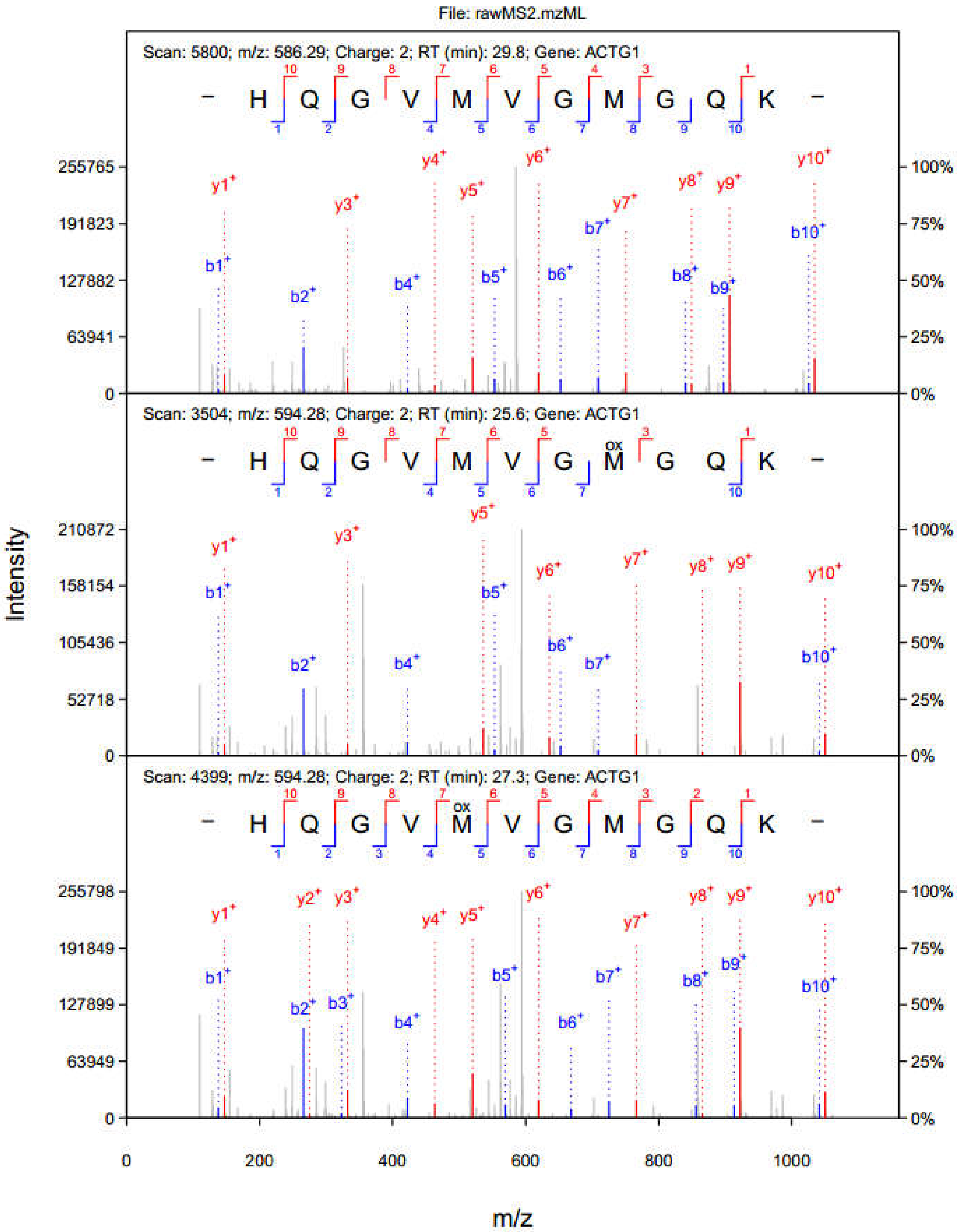

**Figure 6**. The image of the file “rawMS2.mzML_2_HQGVMVGMGQK_align.pdf”.
